# Myeloperoxidase promotes a tumorigenic microenvironment in non-small cell lung cancer

**DOI:** 10.1101/2023.01.28.526014

**Authors:** Paulina Valadez-Cosmes, Kathrin Maitz, Oliver Kindler, Nejra Cosic Mujkanovic, Anna Lueger, Sofia Raftopoulou, Melanie Kienzl, Zala Nikita Mihalic, Ana Santiso, Arailym Sarsembayeva, Luka Brcic, Jörg Lindenmann, Wolfgang Sattler, Akos Heinemann, Rudolf Schicho, Gunther Marsche, A. McGarry Houghton, Julia Kargl

## Abstract

Myeloperoxidase (MPO) is a heme peroxidase that is mainly expressed and secreted by neutrophils. MPO’s role in inflammatory diseases has been highlighted in recent years, but its role in tumor development remains unclear. Therefore, we investigated the role of MPO in non-small cell lung cancer (NSCLC). *In silico* analysis revealed a survival benefit in patients with NSCLC and low MPO expression. Furthermore, a syngeneic tumor model using MPO knockout (KO) mice revealed that mice lacking MPO had lower tumor growth than controls. The reduction in tumor size was accompanied by an increase in lymphoid populations, including natural killer cells and CD8^+^ T cells, suggesting a shift to a more anti-tumorigenic immune environment in MPO-KO mouse tumors. The T cell induced interferon-gamma (IFN-γ) expression was increased in MPO-KO tumors, indicating increased tumoricidal activity. CD8 depletion abolished the previously observed reduction in tumor size in MPO-KO mice, indicating that CD8^+^ T cells play an important role. *In vitro*, T cells treated with MPO showed reduced proliferation and IFN-γ expression. Furthermore, MPO could be internalized into T cells. Heparin pretreatment of T cells blocked MPO binding and internalization into T cells and reversed MPO-induced proliferation reduction. Interestingly, MPO^+^ lymphocytes were found in tumor samples from patients with NSCLC. Our findings suggest that MPO plays an immunosuppressive role in NSCLC.

**One Sentence Summary:** High myeloperoxidase (MPO) expression in non-small cell lung cancer patients is a predictor for adverse outcome and mice lacking MPO showed enhanced anti-tumorigenic leukocyte infiltration, suggesting a pro-tumorigenic role of MPO.

## INTRODUCTION

Lung cancer remains the leading cause of cancer-related mortality worldwide (*1*). Non-small cell lung cancer (NSCLC) accounts for 85% of all lung cancer diagnoses (*2*). Despite recent advances in cancer therapy, the 5-year survival rate for patients with lung cancer remains dismal at around 15% (*3*). The identification of novel therapeutic targets within the tumor microenvironment (TME) is crucial for developing new strategies to treat lung cancer.

Tumor-infiltrating neutrophils and CD8^+^ tumor-infiltrating lymphocytes are two important cell types that shape the TME. Despite their complexities, one of the main functions of CD8^+^ T cells within the TME is to recognize tumor-associated antigens and kill cancer cells (*4*). Indeed, higher CD8^+^ T cell counts have been linked to better outcomes in breast (*5*), colorectal (*6*), and lung (*7, 8*) cancers, among others. On the other hand, neutrophils have been reported to play ambivalent roles in cancer, including pro (*9*–*11*) and anti-tumorigenic (*12, 13*) properties. Growing evidence suggests that tumor-infiltrating neutrophils can act as immunosuppressive cells and manipulate T cell behavior in favor of the tumor (*14*). Infiltrating neutrophils are abundant in NSCLC (*15*). Furthermore, some studies have linked neutrophil infiltration within tumors to poor clinical outcomes (*16*). It has also been demonstrated that neutrophils suppress lymphocyte proliferation and function within the TME (*17, 18*). More recently, some neutrophil cytoplasmic granule components, including myeloperoxidase (MPO), have been proposed to contribute to tumor development (*19*–*21*).

MPO is a heme-containing enzyme that is primarily expressed by neutrophils (*22*) but is also found in small amounts in other myeloid-derived cells such as monocytes (*23*) and macrophages (*24*). MPO is found in neutrophil primary (azurophilic) granules, accounting for 5% of total protein content (*22*). MPO, like other peroxidases, catalyzes the production of many powerful oxidants, most notably the conversion of hydrogen peroxide and chloride ions to hypochlorous acid (HOCl) (*25*). Upon neutrophil activation, MPO is secreted into the extracellular milieu during acute inflammation or in the TME, where it can cause tissue damage at inflammatory sites (*26, 27*). However, little is known about the role of MPO in tumor development.

The purpose of this study was to determine the role of MPO in the progression of lung cancer. In the present study, we show that MPO knockout (KO) mice had smaller tumors than wild-type (WT) littermates, indicating a pro-tumorigenic role for MPO in NSCLC. This was accompanied by the increased accumulation of several lymphoid populations and the local tumoricidal activity of CD8^+^ T cells. *In vitro* analysis revealed that MPO can internalize into human T cells and negatively regulate their proliferation and activity. This study sheds light on a novel role of neutrophil-derived molecules as key players in tumor growth and, for the first time, identifies MPO as an enzyme within the TME with an immunosuppressive role in cancer.

## RESULTS

### MPO expression is associated with shorter survival in patients with NSCLC and positively influences tumor growth in vivo

To determine the significance of MPO in NSCLC, the prognostic value of MPO protein expression in NSCLC was first investigated. Using the Clinical Proteomic Tumor Analysis Consortium (CPTAC) database, the influence of MPO protein expression in primary tumor tissues on survival was investigated using Kaplan–Meier analysis, revealing a benefit for patients with low MPO protein expression [Fig. 1(**A**)]. To further investigate the effect of MPO, patients were divided into MPO-high and -low groups based on the median MPO protein expression, and differentially expressed genes were determined using Welch’s t-test with a cutoff of 0.05 for the adjusted *p*-values [Fig. 1(**B**), table S1]. In the TCGA-lung adenocarcinoma (LUAD) dataset, significantly upregulated genes were used to calculate an MPO signature score [Fig. 1(**B**)]. To validate the use of this signature, the scores of significantly upregulated and downregulated genes were correlated with each other, revealing a strong negative correlation, implying a biological reason driving this expression profile [Fig. S1(**B**) and (**C**)]. The effect of the signature on the progression-free interval (PFI) and overall survival (OS) was evaluated and revealed a higher survival in the low MPO signature group [Fig. 1(**C**) and (**D**)].

**Fig. 1.**
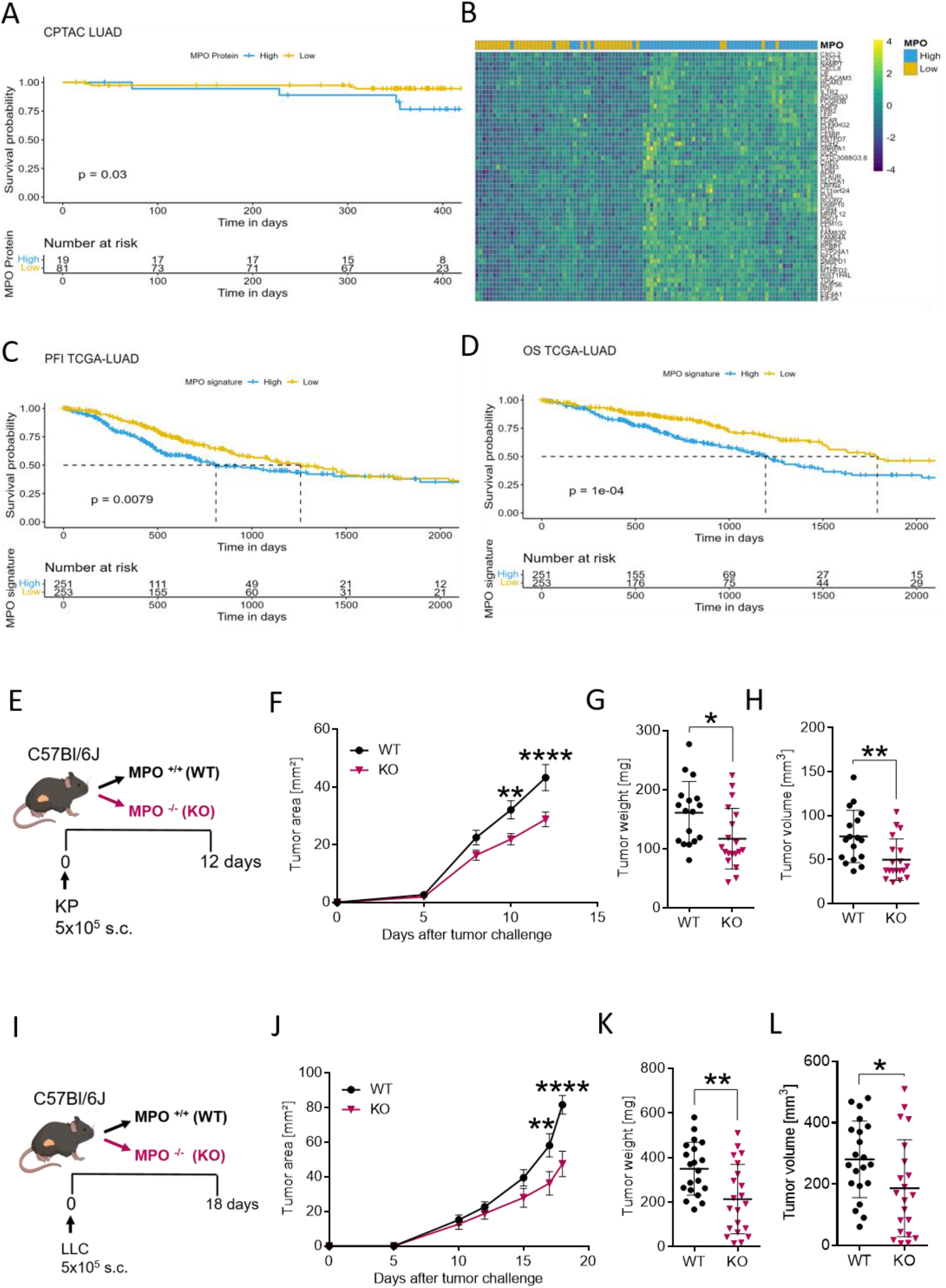
MPO expression is clinically relevant in patients with NSCLC and impacts tumor growth in mice. **(A)** Kaplan–Meier analysis of survival outcome differences in patients with NSCLC and different MPO expression levels. **(B)** Heatmap of differentially expressed genes in MPO-high and -low subgroups. The horizontal axis represents samples, and the vertical axis represents different genes. **(C)** Progression-free survival in TCGA patients divided into MPO-high and -low subgroups. **(D)** Overall survival in TCGA patients stratified by MPO expression. **(E and I)** Wild-type (WT) and MPO knockout (KO) mice were subcutaneously injected with 5 × 10^5^ KP or LLC cells, and tumors were allowed to grow for 12 or 18 days, respectively. **(F and J)** Tumor growth was monitored throughout the experiment. The data were pooled from two independent experiments (*n* = 18–21) and expressed as means ± standard error of the mean. Statistical differences were assessed using a two-way analysis of variance (ANOVA) with Sidak’s post hoc test. ** *p* < 0.01 and **** *p* < 0.0001. **(G, H, K, and L)** Mice were sacrificed, and tumor weight and volume were measured *ex vivo*. The data were pooled from two independent experiments (*n* = 18–21) and expressed as means ± standard deviations. All variables were tested for Gaussian distribution using the Shapiro–Wilk normality test. Statistical differences between WT and KO mouse data with normal distribution were determined using unpaired student’s t-tests with Welch’s correction; otherwise, the Mann– Whitney test was applied (\**p* < 0.05 and \*\**p* < 0.01).

To evaluate how MPO deficiency affects tumor burden in mice, KP cells (isolated from a Kras^LSL-G12D^/p53^fl/fl^ mouse LUAD) or LLC cells (Lewis lung carcinoma) were injected subcutaneously into the flanks of C57BL/6 MPO^+/+^ WT and age-matched MPO-KO (MPO^−/−^) mice [Fig. 1(**E**) and (**I**)]. Tumor growth was monitored for 2 weeks. The tumor growth curves revealed that KP and LLC tumors grew more slowly in MPO-KO animals than in WT animals [Fig. 1(**F**) and (**J**)]. Ex vivo analysis of the weight and volume of KP [Fig. 1(**G**) and (**H**)] and LLC [Fig. 1(**H**) and (**L**)] tumors revealed a 35% and 40% reduction in tumor size in MPO-KO mice, respectively.

Our findings suggest that MPO plays a role in the development of NSCLC and has an impact on patient survival.

### MPO deficiency favors an anti-tumorigenic immune cell profile in NSCLC mouse models

To determine whether the reduced tumor size observed in MPO-KO mice was accompanied by changes in the TME’s immune cell profile, flow cytometry was used to determine the comprehensive profile of infiltrating immune cells and their subtypes in KP and LLC tumors [gating strategies shown in fig. S2(**A**) and (**B**)]. There were no differences in the number of live cells and infiltrating leukocytes (CD45^+^ cells) in KP tumors from MPO and WT mice [Fig. 2(**A**) and (**B**)]. MPO-KO mice had more monocytes and eosinophils than WT mice [Fig. 2(**C**)], but no changes in the other myeloid cell populations, including neutrophils, macrophages, dendritic cells (DCs), myeloid dendritic cells (mDCs) and plasmacytoid dendritic cells (panDCs) [Fig. 2(**C**)]. Additionally, we observed a significant shift in lymphoid cell populations in MPO-KO tumors compared to WT mice, with γδT cells, natural killer (NK) cells, NKT cells, T cells (CD3^+^), and CD8^+^ T cells being increased in MPO-KO mice [Fig. 2(**D**) and (**E**)]. Interestingly, the relative abundance of memory CD4^+^ and CD8^+^ T cells, as well as effector CD8^+^ T cells, was higher in MPO-KO mice than in WT mice [Fig. 2(**F**) and (**G**)]. The expression of the inhibitory checkpoint receptor PD-1 on tumor-infiltrating CD4^+^ and CD8^+^ T cells was increased in tumors from MPO-KO mice, suggesting increased immune activity of these cells [Fig. 2(**H**) and (**I**)]. MPO-KO mice had more live and CD45^+^ cells in their LLC tumors than WT mice [fig. S3(**A**) and (**B**)]. Furthermore, B cells, NK cells, NKT cells, CD3^+^ T cells, CD4^+^ T cells, and CD8^+^ T cells were also increased [Fig. S3(**C**) and (**D**)]. The relative abundance of naïve and effector CD8^+^ T cells was higher in MPO-KO mice [Fig. S3(**F**)]. The expression of the inhibitory checkpoint receptor PD-1 on tumor-infiltrating CD8^+^ T cells was also increased in MPO-KO mice [Fig. S3(**H**)].

**Fig. 2.**
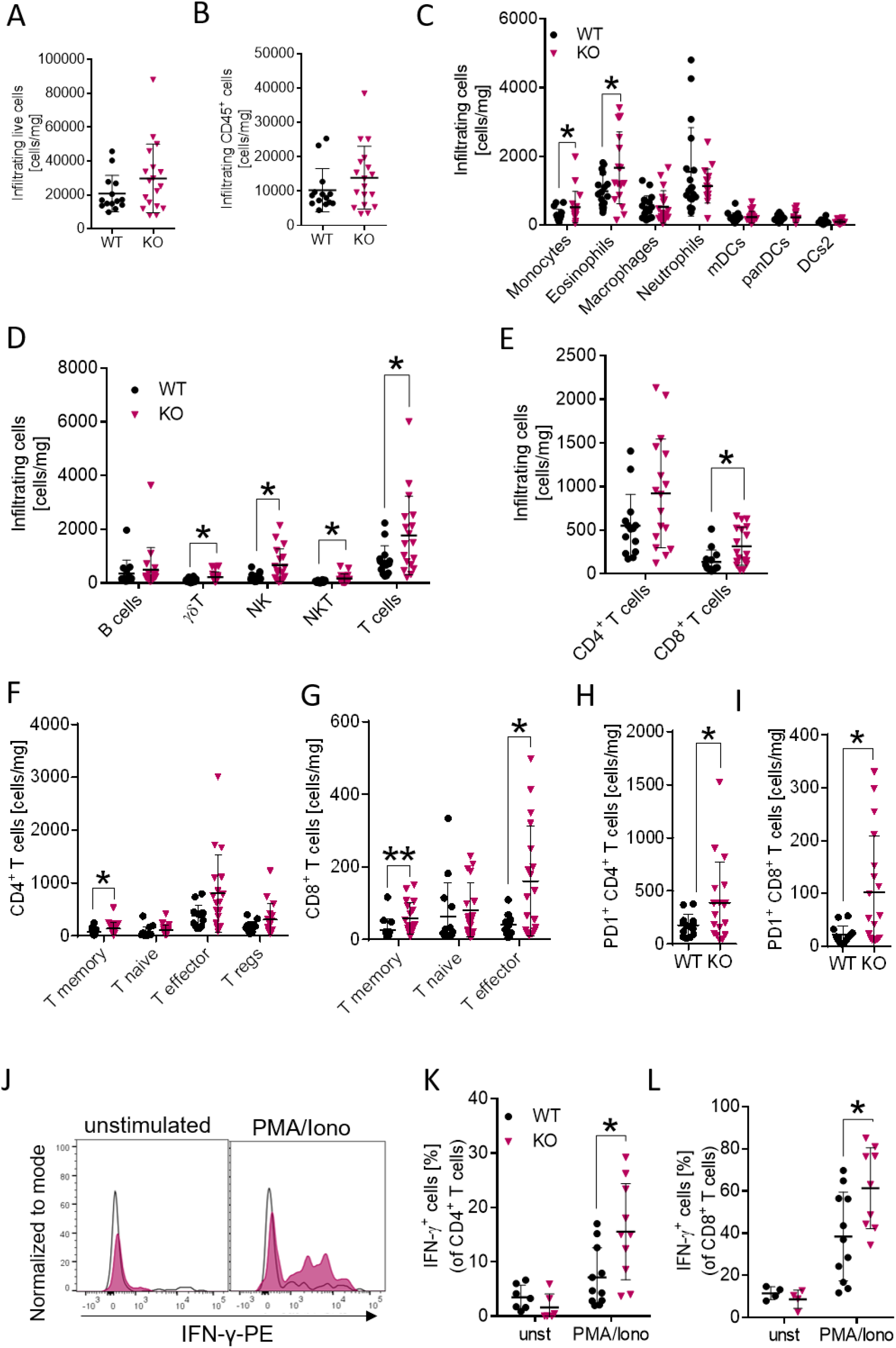
MPO deletion favors lymphocyte infiltration in an engraftment lung cancer model. **(A–I)** Flow cytometric analysis of single-cell suspensions of KP tumors from MPO wild-type (WT) and knockout (KO) mice. The data were pooled from two independent experiments (*n* = 18– 20) and expressed as means ± standard deviations. All variables were tested for Gaussian distribution using the Shapiro–Wilk normality test. Statistical differences between WT and KO mouse data with normal distribution were determined using unpaired student’s t-tests with Welch’s correction; otherwise, the Mann–Whitney test was applied (\**p* < 0.05 and \*\**p* < 0.01). **(J)** Representative histograms of IFN-γ expression in tumor-infiltrating CD8^+^ T cells before (unstimulated) and after PMA/Iono stimulation. **(K and L)** Quantitative analysis of tumor-infiltrating CD4^+^ and CD8^+^ T cells revealed increased IFN-γ expression in MPO-KO mice after ex vivo PMA/Iono stimulation. DCs, dendritic cells; mDCs, myeloid dendritic cells; panDC, plasmacytoid DCs; iono, ionomycin; PMA, phorbol 12-myristate 13-acetate/ionomycin; NK, natural killer cells; NKT, natural killer T cells

When WT mice were compared to MPO-KO mice, no differences were observed in any of the immune populations in the spleens of nontumor-bearing [Fig. S4(**A**)–(**E**)] or tumor-bearing mice [Fig. S6(**A**)–(**E**)]. The thymus of tumor-free mice revealed a slight increase in the CD8^+^ single-positive thymocyte subpopulation (CD8^+^ SP) in MPO-KO mice compared to WT mice [Fig. S5(**C**)]. When compared to the WT, cells of the differentiation stage DN1 were reduced in the MPO-KO while cells of the differentiation stage DN4 were increased [Fig. S5(**D**)]. There were no differences in the other thymocyte populations [Fig. S5(**A**)–(**D**)].

To test whether T cells were more activated in MPO-KO mice, tumor-infiltrating CD8^+^ and CD4^+^ T cells from MPO-KO and WT mice were stimulated ex vivo with phorbol 12-myristate 13-acetate/ionomycin (PMA/Iono) and their activity was assessed using flow cytometry. MPO-KO tumors expressed higher IFN-γ levels on CD8^+^ and CD4^+^ T cells than WT mice [Fig. 2(**J**)– (**L**)], indicating local activation and enhanced tumoricidal activity of T cells in MPO-KO mice. Our findings, therefore, suggest that MPO depletion increases lymphocyte infiltration into the tumors.

### Reduction of tumor size in MPO-KO mice is dependent on CD8^+^ T cells

Given the differences in CD8^+^ T cell counts, KP tumor-bearing mice were injected intraperitoneally with anti-CD8 antibodies or control IgG isotypes to explore whether the reduction in tumor size in the MPO-KO mice is dependent on the presence of CD8^+^ T cells [Fig. 3(**A**)]. At the endpoint, tumors were harvested, weighed, measured, and processed for flow cytometry. The CD8^+^ T cell pool was depleted by 90% on average (Fig. S7). In the isotype-treated mice, tumor weight and volume were reduced in the MPO-KO group vs the WT [Fig. 3(**B**) and (**C**)]. Interestingly, in the anti-CD8 antibody-treated mice, no differences in tumor weight and volume were observed between the WT and MPO-KO groups [Fig. 3(**B**) and (**C**)]. Flow cytometric analysis of tumor single-cell suspensions revealed no differences in CD45^+^ infiltration between WT and MPO-KO mice in both the isotype and anti-CD8 antibody groups [Fig. 3(**D**)]. As previously found, CD8^+^ T cells were found to be higher in isotype-treated MPO-KO mice than in WT mice [Fig. 3(**E**)]. Our findings suggest that CD8^+^ T cells are required for MPO-KO-dependent tumor growth reduction.

**Fig. 3.**
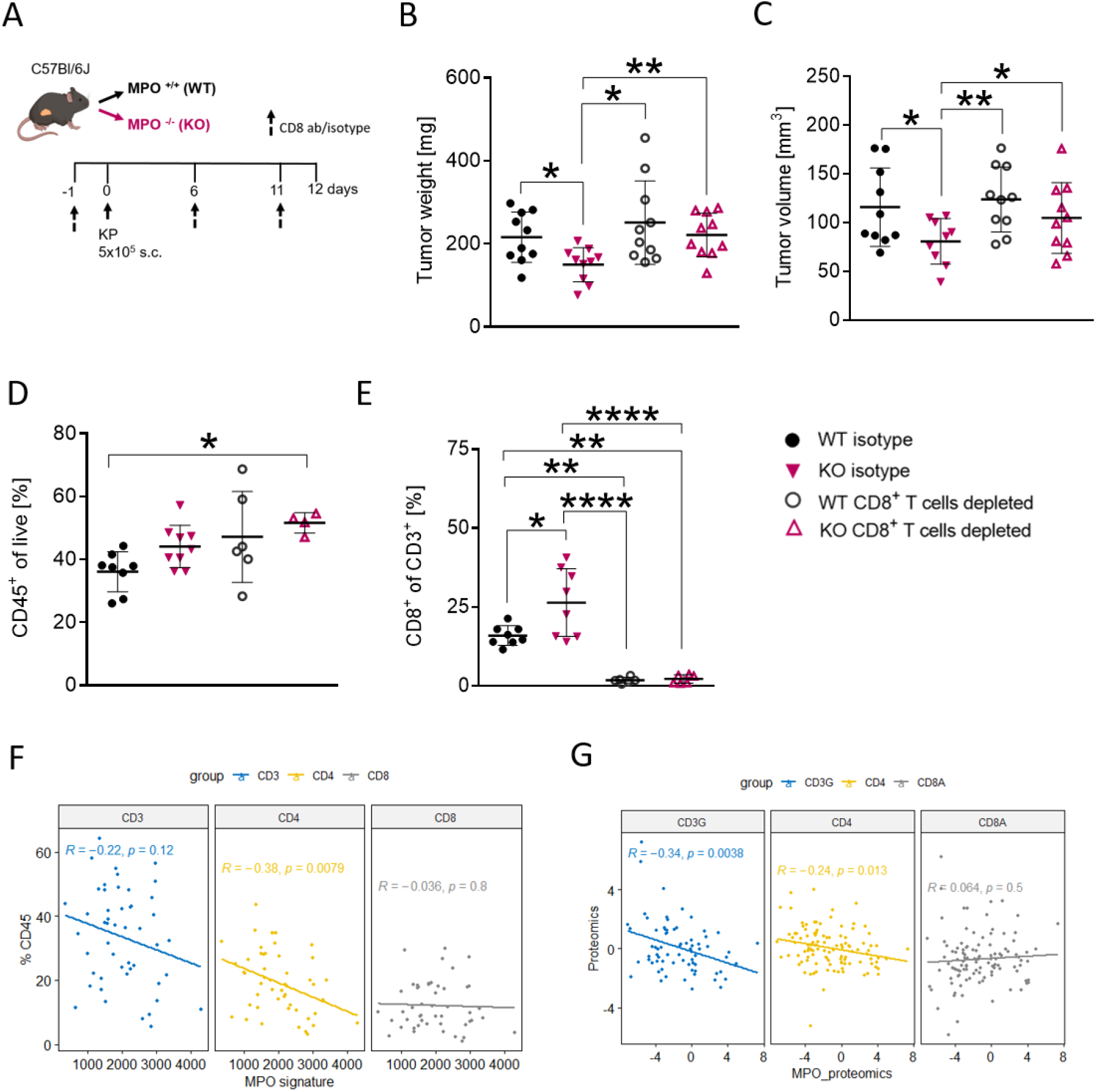
CD8+ T cells are necessary to reduce tumor growth in MPO-KO mice. **(A)** Schematic representation of the subcutaneous tumor model used in WT and MPO-KO mice. Anti-CD8 antibody (500 µg per mouse) was given intraperitoneally on Days −1 (1 day prior to tumor cell injection) and 6, and 250 µg of anti-CD8 antibody were given intraperitoneally on Day 11. **(B and C)** One day after the last anti-CD8 antibody injection, mice were sacrificed, and tumor weight and volume were measured ex vivo. The data were gained from one independent experiment (*n* = 10) and expressed as means ± standard deviations. All variables were tested for Gaussian distribution using the Shapiro–Wilk normality test. Statistical differences between WT and KO mouse data with normal distribution were determined using unpaired student’s t-test with Welch’s correction (* *p* < 0.05, ** *p* < 0.01, and **** *p* < 0.0001). **(D and E)** Flow cytometric analysis of single-cell suspensions from tumors in WT and MPO-KO mice. The data were gained from one independent experiment (*n* = 6–10) and expressed as means ± standard deviations. Statistical differences between WT and KO mouse data were assessed using one-way analysis of variance (ANOVA) with Tukey’s multiple comparisons test (\**p* < 0.05, \*\**p* < 0.01, and \*\*\*\**p* < 0.0001). **(F)** Scatter plots depicting Pearson’s correlation between the MPO signature and T cell subsets in the ICP3 cohort. **(G)** Scatter plots depicting Pearson’s correlation between the MPO signature and T cell subsets in the CPTAC-LUAD cohort.

In the ICP3 cohort, and the CPTAC-LUAD cohort, Pearson’s correlation between the MPO signature and T cells revealed a negative correlation between MPO protein expression and CD3^+^ T cells and CD4^+^ T cells [Fig. 3(**F**) and (**G**)]. Interestingly, no correlation was found between MPO protein expression and CXCL9, CXCL10 and CCL5 cytokine gene expression in the CPTAC-LUAD cohort [Fig. S1(**D**)].

### MPO decreases T cell proliferation and function and internalizes into T cells in vitro

Since tumors in MPO-KO mice had higher T cell infiltration and tumor reduction in MPO-KO mice was dependent on CD8^+^ T cells, the underlying mechanisms by which MPO could affect T cell behavior were investigated *in vitro*. T cells were isolated from healthy donors’ PBMCs, stained with the proliferation dye eFlour™ 450, and incubated with MPO for 72 h. CD4^+^ and CD8^+^ T cell proliferation was significantly reduced by about 20% after 72-h incubation with MPO [0, 5, 10, 20 μg/ml; Fig. 4(**A**)–(**C**)]. To investigate the effect of MPO on T cell activation, human CD8^+^ and CD4^+^ T cells were treated with MPO (0, 5, 10, 20 μg/ml) for 24 h, stimulated with PMA/Iono, and assessed for activity as measured by their ability to produce IFN-γ using flow cytometry. MPO-treated CD4^+^ and CD8^+^ T cells showed decreased IFN-γ expression levels compared to vehicle-treated cells [Fig. 4(**D**) and (**E**)], indicating a reduced T cell activation. These findings imply that MPO has a direct impact on T cell function by reducing proliferation and activation.

**Fig. 4.**
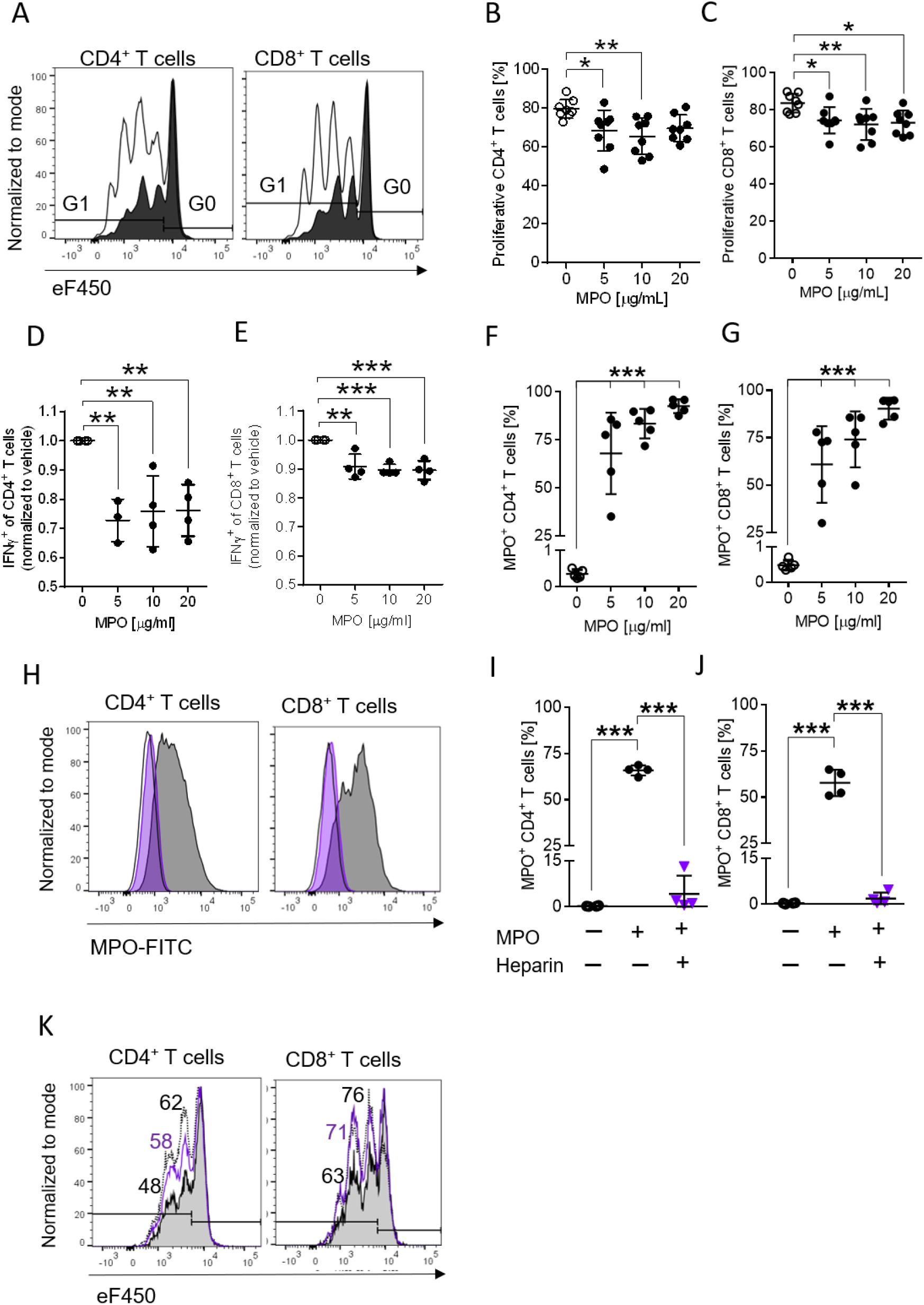
MPO binds to T cells and inhibits their proliferation and function *in vitro*. PBMCs were isolated from healthy donors, and the T cell fraction was enriched using negative selection. **(A–C)** 2.5 × 10^5^ T cells were labeled with the proliferation dye eFlourTM 450, treated with different MPO concentrations, and cultured for 72 h. Cells were stained for CD4 and CD8, and cell proliferation was evaluated as an increase in the percentage of dividing cells (G1). **(D and E)** 2.5 × 10^5^ T cells were incubated with MPO for 24 h. To stimulate IFN-γ production, cells were treated with ionomycin (1 μg/ml) and PMA (100 ng/ml). In addition, Golgi stop (1,5 μg/ml) was used to prevent IFN-γ secretion. After incubation, cells were stained for CD4 and CD8, and IFNγ expression was evaluated using flow cytometry. **(F and G)** T cells were treated with MPO (0, 5, 10, and 20 μg/mL) for 2 h, and intracellular/nuclear flow cytometric staining was performed. **(H– J)** T cells were pretreated with heparin for 45 min, washed, and then treated with MPO (10 μg/mL) for 2 h. **(A-J)** The data were pooled from four to eight independent experiments and expressed as means ± standard deviations. Statistical differences were determined using one-way ANOVA with Sidak’s post hoc test (\**p* < 0.05, \*\**p* < 0.01 and \*\*\**p* < 0.001). **(K)** T cells were pretreated with heparin, and the proliferation was assessed in CD4^+^ and CD8^+^ T cells. Numbers represent the percentage of proliferative cells in the vehicle (dotted line), MPO-treated (black line), and heparin-MPO-treated (purple line) cells (*n* = 1).

Although MPO is not expressed by T cells, several studies have reported its ability to internalize into endothelial and epithelial cells (*28, 29*). Accordingly, to investigate MPO’s ability to bind and internalize into T cells, cells were incubated for 2 h in the presence of MPO (0, 5, 10, and 20 μg/mL), and MPO binding and internalization were assessed using flow cytometry. After 2 h of treatment, more than 50% of CD4+ and CD8+ T cells were MPO-positive [Fig. 4(**F**) and (**G**)]. It was previously shown that MPO internalization is dependent on cell surface glycosaminoglycan interactions, and soluble glycosaminoglycans such as heparin can block MPO cellular uptake (*28*). Heparin pretreatment of T cells for 45 min completely blocked MPO binding to T cells [Fig. 4(**H**)–(**J**)]. Interestingly, heparin pretreatment of T cells reversed MPO’s effect on T cell proliferation [Fig. 4(**K**)], implying that MPO binding and internalization to T cells are required for its functional role.

### MPO is found in the lymphocytes of tumor samples from patients with NSCLC

To investigate whether our findings about MPO binding and internalization into T cells were replicated in clinical samples, we stained human LUAD tissue sections and assessed NSCLC tissues for the presence of MPO in lymphocytes by flow cytometry. Immunohistochemistry staining of LUAD tissues revealed the presence of MPO in lymphocytes, which is consistent with our *in vitro* findings [Fig. 5(**A**)]. Accordingly, flow cytometry revealed a lymphocyte MPO^+^ population in several NSCLC samples [Fig. 5, (**B**)–(**E**)]. Interestingly, a positive correlation was found between the percentages of infiltrated neutrophils [defined as CD163^−^ CD66b^+^, Fig. S2(**C**)], the main source of MPO in NSCLC, and the percentages of MPO^+^ lymphocytes in NSCLC samples [Fig. 5(**F**)].

**Fig. 5.**
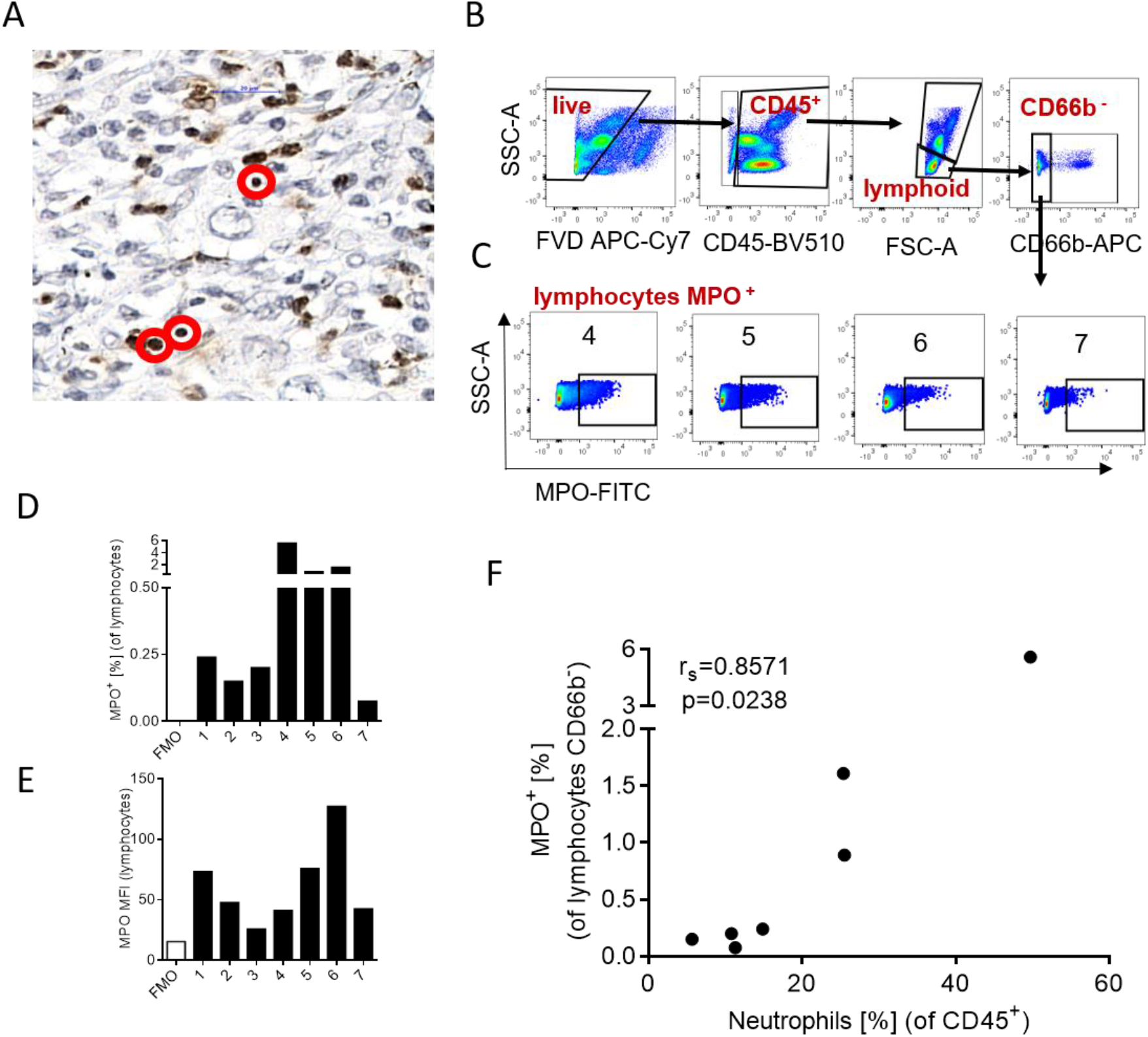
MPO is found in lymphocytes from NSCLC tumor samples. **(A)** MPO staining in a variety of inflammatory cells, including lymphocytes (circle) (digital magnification: ×80). **(B)** Representative flow cytometric dot plots demonstrating the gating strategy used to identify MPO^+^ signal in lymphocytes from patients with NSCLC. The initial gate is on live (FVD) and CD45^+^ cells. The lymphoid population (T, B, and NK cells) was gated based on its forward and side scatter properties. The CD66b marker was used to exclude myeloid cells within this gate **(C and D)** MPO^+^ lymphocytes in patients with NSCLC.**(E)** MPO median fluorescence intensity. **(F)** Correlations between MPO+ lymphocytes and neutrophil content in seven patients with NSCLC. The correlation was determined using Spearman’s correlation coefficient rho (rs).

## DISCUSSION

The importance of the TME as a niche that not only contains different cell types that modulate tumor growth but also delivers vital factors that support the escape from host immune surveillance and thus promote cancer development is well described (*30*). In this context, neutrophils and neutrophil-derived molecules have been identified as important players in NSCLC development (*15*). Molecules like neutrophil elastase have been shown to play a pro-tumorigenic role in NSCLC (*20*). Accordingly, we aimed to determine whether another important neutrophil-derived granular enzyme, MPO, contributes to NSCLC carcinogenesis.

MPO is a heme peroxidase mainly expressed by neutrophils (*22*) but is also found in small amounts in monocytes and macrophages (*27*). MPO is normally stored in neutrophil primary granules, but it can be rapidly released into the extracellular milieu in response to neutrophil activation, apoptosis, and necrosis. MPO’s physiological roles have traditionally been described in the context of an innate immune response, in which this enzyme is responsible for pathogen killing once neutrophils arrive at the site of infection (*31*). Recently, emerging roles of the enzyme have been identified, particularly in the context of cardiovascular disease and atherosclerosis, where MPO has been linked to both the initiation and progression of cardiac pathologies (*32, 33*). Since then, special interest has emerged in studying the role of MPO in other inflammatory diseases such as rheumatoid arthritis, cutaneous and pulmonary inflammation, and cancer (*34*).

In the present study, patients with NSCLC and low MPO protein expression had a better survival prognosis than patients with high MPO protein expression. Because MPO gene expression is restricted to neutrophil precursors in the bone marrow (*35*), MPO expression in solid tumor RNA datasets does not correlate with MPO protein expression [Fig. S1(**A**)]. To address this limitation, the MPO signature was calculated to exploit MPO’s role in publicly available gene expression data. The MPO-low signature was linked to better survival (PFI and OS). Consistent with our findings, higher MPO expression has been linked to increased oxidative stress and higher susceptibility to lung (*36*), ovary (*37*), and breast (*38*) cancer, among others. Furthermore, some MPO-derived oxidants can induce carcinogenesis by inducing DNA damage (*39*).

Using a syngeneic lung tumor model, we observed that tumors grown in MPO-deficient mice were smaller than those grown in WT mice. Our findings are consistent with those of Rymaszewski et al., who found that MPO-KO mice had smaller tumors than control mice (*40*). In the same study, pharmacologic MPO inhibition had no effect on tumor size when compared to controls (*40*). However, early administration of an MPO inhibitor in an inflammation-driven lung tumor model remarkably reduced lung tumor multiplicity, suggesting an immunoregulatory effect (*40*).

Interestingly, our study discovered that the immune cell landscape of MPO-KO tumors had shifted to a more anti-tumorigenic profile. Significant increases in lymphoid populations, including NK, NKT, γδ T cells, and CD3^+^CD8^+^ T cells were observed in MPO-KO mouse tumors compared to their WT counterparts. In line with our findings, increased immune infiltration of cytotoxic T cells such as CD3^+^CD8^+^ T cells and NK cells has been linked to a better prognosis for lung cancer (*41, 42*). A more in-depth investigation of CD4^+^ and CD8^+^ T cells revealed that they were also more active with higher IFN-γ levels. Previous reports found that patients with NSCLC had a lower number of cytotoxic T cells as well as IFN-γ expression (*43, 44*). Hence, MPO deficiency may contribute to tumor shrinkage by improving the local immune microenvironment and boosting the activity of infiltrated lymphocytes. Further analysis of CD8^+^ T cell subpopulations revealed an increase in effector and memory CD8^+^ T cells, which is consistent with previous studies that found a reduction in T effector cell infiltration in patients with NSCLC (*44*). Another interesting finding was that PD-1 expression was higher in CD4^+^ and CD8^+^ lymphocytes of MPO-KO mice, implying strong T cell activity against tumor antigens as well as a possible predictor of anti-PD-1 therapy response (*45, 46*). Interestingly, another study found that using an MPO inhibitor in combination with PD-1 checkpoint blockade reduced tumor progression (*47*) pointing to MPO as a potential therapeutic target to improve current cancer immunotherapies. To the best of our knowledge, this is the first study to link MPO deficiency to a shift in immune populations infiltrating tumors.

Cytotoxic CD8^+^ T cells play an important role in the anticancer immune response (*4*). To determine whether the increased number of CD8^+^ T cells was linked to a reduced tumor burden in MPO-KO mice, anti-CD8 antibodies were used to deplete CD8^+^ T cells. Remarkably, no differences in tumor weight or volume were observed between the WT and MPO-KO groups when CD8^+^ T cells were abolished. Thus, the impact on the CD8^+^ T cell population may represent an important mechanism against tumorigenesis caused by MPO deficiency in the TME. In line with our *in vivo* findings, we found that MPO protein expression was negatively correlated with CD3^+^ and CD4^+^ T cells in NSCLC human samples. However, the correlation between MPO protein expression and CD8^+^ T cells was found to be insignificant.

Tumor cells produce cytokines and chemokines that attract leukocytes, including T lymphocytes, to the tumor site. Interestingly, *in silico* data revealed no correlation between MPO protein expression and cytokine content in human NSCLC samples [Fig. S1(**D**)]. On the other hand, human T cells treated with MPO had reduced proliferation and IFN-γ production *in vitro*. These findings suggest that MPO has a direct impact on T cell behavior. Other studies have shown that after antigen injection, neutrophils rapidly infiltrate draining lymph nodes where MPO is released (*48*). The deposited MPO suppresses DC function and migration, resulting in a decrease in CD4^+^ T cell responses, including T cell activation, proliferation, and differentiation into Th1 and Th17 effectors (*48*). Accordingly, MPO-driven lipid peroxidation has recently been described as a mechanism by which MPO impairs antigen cross-presentation by DCs in tumors (*47*).

MPO is a highly cationic molecule that rapidly binds to negatively charged structures such as bacterial surfaces, extracellular matrix components, and cellular membranes, including those of endothelial cells and neutrophils themselves (*28, 49*). Moreover, extracellular MPO uptake has been reported in neutrophils and macrophages via mechanisms involving CD11b/CD18 integrins and the mannose receptor, respectively (*50, 51*). MPO uptake has previously been demonstrated in endothelial cells, and a functional impact of MPO after binding and internalization has also been reported (*51*). Accordingly, we found that MPO could bind to and internalize into T cells, and that heparin pretreatment completely blocked the presence of MPO in T cells. This is consistent with another study that found the same effect in endothelial cells (*28*). Furthermore, we found MPO^+^ lymphocytes in NSCLC samples, and MPO^+^ lymphocytes were positively correlated with neutrophil abundance. Remarkably, the reduction in T cell proliferation observed after MPO treatment was reversed when T cells were pretreated with heparin, implying that MPO binding and/or internalization into T cells is required for its functional role. More research into the mechanism by which MPO impairs T cells function is required.

In conclusion, our findings suggest that MPO in the NSCLC TME may act as an immunosuppressive molecule, impairing T cell behavior and thus promoting tumor growth.

## MATERIALS AND METHODS

### Bioinformatics

#### CPTAC cohort

CPTAC LUAD protein, transcript, clinical, and survival data were accessed using the cptac python package (https://pypi.org/project/cptac/). Patients were divided into groups based on the median MPO protein expression, and *Welch’s t*-test was used to identify differentially expressed genes. A critical *p*-value of 0.05 was chosen, and Bonferroni correction was used as an adjustment method. Significantly upregulated genes were plotted on a heatmap using the R package pheatmap, and this list of genes was further termed the “MPO signature.” Significantly downregulated genes were termed the “control signature” and were used as a control in other datasets to assess the plausibility of the MPO signature. MPO protein and transcript expressions were correlated using Pearson’s correlation and plotted using the R package ggpubr. The survival curve was created using the R package survminer. The cutpoint was determined using the surv_cutpoint function of this package.

#### The Cancer Genome Atlas (TCGA)-LUAD cohort

Transcriptomic (TPM), clinical, and survival data were accessed via the pan-cancer atlas hub on the UCSC Xena browser. The MPO signature was calculated using the z-score method in the R package hacksig. The Kaplan–Meier analysis was carried out using the R package survminer, and the median expression of the MPO signature was used as a cutpoint.

#### ICP3 cohort

The MPO signature was calculated using the z-score method in the R package hacksig and correlated with T cell content (% CD45+) derived from flow cytometry (*15*).

#### MPO signature plausibility check

In datasets where the MPO signature was used, MPO signature and control scores were scaled using R’s scale function and correlated using Pearson’s correlation. In the ICP3 cohort, the neutrophil content determined by flow cytometry was correlated with the scaled MPO signature score. In the TCGA-LUAD cohort, the neutrophil content obtained from the Kassandra website (https://science.bostongene.com/kassandra/downloads, 10.1016/j.ccell.2022.07.006) was correlated with the scaled MPO signature score.

Software: Python 3.10.6, R 4.2.1, Platform: x86_64-conda-linux-gnu (64-bit), operating system: Debian GNU/Linux 11 (bullseye).

### Animal studies and cell culture

All animal experiments were performed at the animal facilities of the Medical University of Graz. The experimental protocols were approved by the Austrian Federal Ministry of Science and Research (BMWF-66.010/0041-V/3b/2018). MPO^*−/−*^ mice (B6.129X1-Mpo^tm1Lus^/J) were obtained from the Jackson Laboratory (Bar Harbor, ME, USA) and backcrossed to the C57BL/6J strain. Genotyping was performed from ear tags using standard PCR protocols. The murine KP cell line (generously provided by Dr. A. McGarry Houghton, FredHutch, Seattle) was isolated from a LUAD of a Kras^LSL-G12D^ / p53^fl/fl^ mouse at the Fred Hutchinson Cancer Center (Seattle, WA, USA) after intratracheal administration of adenoviral Cre recombinase, as previously described (*52*). The murine LLC cell line (ATCC® CRL-1642™) as well as the human LUAD cell line A549 (ATCC® CCL-185™) were supplied by ATTC. KP and LLC cells were cultured in Dulbecco’s Modified Eagle Medium supplemented with 10% fetal bovine serum (FBS, Life Technologies; catalog nos. 21875–091 and 10270106, respectively) and 1% penicillin/streptomycin (P/S, PAA Laboratories; catalog no. P06-07100) and incubated at 37 °C and 5% CO_2_ in a humidified atmosphere.

### Murine tumor models

For the heterotopic lung tumor engraftment model, KP or LLC cells were subcutaneously injected under inhaled isoflurane anesthesia. A total of 5 × 10^5^ KP or LLC cells suspended in 450 µl Dulbecco’s phosphate-buffered saline (PBS, Gibco) were injected subcutaneously into the flanks of MPO-KO and WT mice aged 6–14 weeks. To deplete CD8 T cells *in vivo*, tumor-bearing MPO-KO mice and WT littermates were injected intraperitoneally with 500 µg of rat monoclonal antibody to mouse CD8α (clone YTS 169.4) (BioxCell; catalog no. BE0117) on Days −1 (1 day prior to tumor cell injection) and 6, and with 250 µg on Day 11, or with rat IgG2b isotype control antibody (Clone LTF-2, BioxCell; catalog no. BE0090). When tumors became palpable (after about 5 days), mice were shaved, and tumor growth was monitored and measured (length and width) every other day using a digital caliper throughout the experiment. Mice were sacrificed on Day 12 (KP cells) or Day 18 (LLC cells), and tumors were harvested, weighted, and measured with a digital caliper *ex vivo*. Tumor volume was calculated using the formula: *V* = *lenght* × *width* × *hight* π/6)(*52*). Tumors were kept in Roswell Park Memorial Institute (RPMI) on ice until further use or were immediately fixed in 10% RotiR-Histofix.

### Human NSCLC tissue samples

Patients with NSCLC Stages IA-IIIB were recruited from the Departments of Internal Medicine, Oncology, and Surgery, Division of Thoracic Surgery, Medical University of Graz (Graz, Austria). Informed consent was obtained from all participants. The study adhered to the Helsinki Declaration and was approved by the Ethics Committee of the Medical University of Graz (MUG-EK-number: 30-105 ex17/18).

### Single-cell suspensions

Single-cell suspensions from mouse and human tumors were prepared as previously described (*52*). Tissue was minced with surgical scissors and digested with collagenase IV (CLS-1; 4.5 U/ml; Worthington) and DNase I (160 mU/mL, Worthington; LS002006) in RPMI for 20 min at 37 °C while rotating at 1000 rpm. Tissue digests were passed through a 100-μm strainer, enzymes were inhibited by adding staining buffer (SB, PBS+2 % FBS), and centrifuged for 5 min at 500 ×g (4 °C). The pellet was resuspended in red blood cell lysis buffer (BioLegend, catalog no. 420301, all human samples) and incubated for 3 min on ice with periodic shaking. The lysis process was neutralized by adding 4 volumes of PBS. The cells were then passed through a 40 μm strainer into a new 50-mL tube and washed in SB. Subsequently, cells were washed twice in PBS, counted using the EVE automated cell counter (NanoEnTek), and stained with surface and intracellular antigens.

### Flow cytometry

#### Mouse

To exclude dead cells, single-cell suspensions were incubated for 20 min at 4 °C in the dark in Fixable Viability Dye (FVD) eFluorTM 780 (eBioscience; catalog no. 65-0865-14). After adding 1 μg TruStain FcX™ (Biolegend; catalog no. 101320) for 10 min, immunostaining was performed for 30 min at 4 °C (protected from light) with the following antibodies (table S1): CD45-AF700 (catalog no. 103128), CD45-BV785 (catalog no. 103149), Ly6C-APC (catalog no. 128015), Ly6G-PE/Dazzle594 (catalog no. 127648), CD11c-BV605 (catalog no. 117334), CD8-PerCPCy5.5 (catalog no. 100734), PD1-APC (catalog no. 135210), CD25-BV785 (catalog no. 102051) CD62L-BV605 (catalog no. 104438), NKp46-BV510 (catalog no. 137623), CD19-FITC (catalog no. 115506), PDL1-PECy7 (catalog no. 124313), CD206-FITC (catalog no. 141703), MHCII-PerCP-Cy5.5 (catalog no. 107625), CCR3-BV421 (catalog no. 144517), CD103-BV510 (catalog no. 121423) (all antibodies from Biolegend), and CD11b-BUV737 (catalog no. 612801), F4/80-BUV395 (catalog no. 565614), Siglec-F-PE (catalog no. 562068), CD3-BUV395 (catalog no. 563565), CD4-BUV496 (catalog no. 564667), CD44-BUV737 (catalog no. 612799), gdTCR-PECF594 (catalog no. 563532) (all antibodies from BD Biosciences), and FoxP3-PE (eBio, catalog no. 12-5773-82). For nuclear antigen staining, cells were permeabilized with Transcription Factor Buffer Set (BD Biosciences, catalog no. 562574) prior to staining procedures. To detect changes in interferon-gamma (IFN-γ), single-cell suspensions of tumors or spleens (2 × 10^6^ cells per well) were seeded into 96-well U-bottomed plates with RPMI containing 10% FBS, 1% P/S, and GolgiStop (1.5 μl/ml, BD Biosciences), and incubated for 4 h at 37 °C. During that time, they were either stimulated with phorbol 12-myristate 13-acetate/ionomycin (PMA, 100 ng/ml, Sigma-Aldrich) and ionomycin (1 μg/ml, Sigma-Aldrich) or left unstimulated. Subsequently, surface staining was performed with CD45-FITC (catalog no. 103108), CD3-BV421 (catalog no. 100336), CD4-PE-Cy7 (catalog no. 100528), and CD8-PerCP5.5 (catalog no. 100734), followed by intracellular labeling (BD Cytofix/CytopermTM Kit) with IFN-γ-PE (catalog no. 505808) and granzyme-B-AF647 (catalog no. 515405). Stained cells were washed, fixed with IC fixation buffer (ThermoFisher Scientific, catalog no. 00-8222-49) for 10 min at 4 °C, and stored at 4 °C until analysis. The samples were analyzed on a BD LSRFortessa™ or a BD Canto™ flow cytometer with FACSDiva software (BD Biosciences, Franklin Lakes, NJ, USA) and over 2 × 10^5^ events per sample were recorded. The data were compensated and analyzed using FlowJo software (TreeStar, Ashland, OR, USA) and gates were defined using fluorescence-minus-one samples [Fig. S2(A)– (**D**)].

#### Human

The staining was performed as described above. In brief, FVD eFluor™ 780 (FVD, eBioscience) (30 min, 4 °C) was used to exclude dead cells. Cells were preincubated in SB with humanTruStain FcX blocking solution (FcBlock, Biolegend) at 4 °C for 10 min. Subsequently, cells were stained in SB for 20 min at 4 °C with CD45-BV510, CD163-BV605, CD66b-APC, and MPO-FITC antibodies (all from Biolegend). Cells were centrifuged (500 ×g, 5 min, 4 °C), resuspended in 200-μL SB, and washed again with SB. Cells were fixed in IC fixation buffer (eBioscience) for 10 min at 4 °C, centrifuged, and resuspended in 100-μL SB. The samples were measured on a BD LSR II Fortessa (BD Biosciences).

### Preparation of human peripheral blood leukocytes

Blood was drawn from healthy volunteers in accordance with a protocol approved by the Ethics Committee of the Medical University of Graz (EK-numbers: 17-291 ex 05/06), as previously described (*53*). Blood (35 ml) was collected in a 50-mL sodium citrate-containing tube. Platelet-rich plasma was separated by centrifugation at 300 ×g for 20 min, and erythrocytes by dextran sedimentation. Dextran T-500 (Sigma-Aldrich; 6 mL, 6% in saline) was added to the cells and filled up to 50 mL with 0.9% saline. Dextran-crosslinked erythrocytes were allowed to sediment for 30 min, and the upper phase containing nonsedimented leukocytes was layered on top of 15 mL Histopaque (Sigma-Aldrich). High-density polymorphonuclear leukocytes, including neutrophils and eosinophils, were separated from less dense peripheral blood mononuclear cells (PBMCs), which included basophils, monocytes, lymphocytes, and dendritic cells, by centrifugation at 350 ×g for 20 min. The PBMCs were isolated from the interphase and washed in a Ca²^+^ and Mg²^+^-free assay buffer.

### T cell isolation

T cells were enriched using the Easy Sep Human T cell Isolation Kit (Stemcell Technologies) according to the manufacturer’s instructions. Briefly, 50 × 10^6^ PBMCs/mL were resuspended in SB and PBS containing 2% FBS and 1 mM EDTA, and then an isolation cocktail solution was added to the PBMCs and incubated for 5 min at room temperature. After incubation, RapidSphere beads were added, and the tube was placed in a magnet for 3 min at room temperature. The enriched sample was collected and washed with PBS. Live cells were counted using a Neubauer chamber and trypan blue. T cells were cultured in prewarmed RPMI media containing 1% P/S and were treated with different MPO (Elastin Products, catalog no. MY862) concentrations at various time points.

### T cell proliferation

T cells were labeled with eFluorTM 450 (eF450, 10 μM in PBS; 10 mio cells/mL; Thermo Fisher) for 10 min. Cells were washed twice with SB and resuspended in T cell proliferation medium (1% P/S, 50-μM β-mercaptoethanol, 2-mM L-Glutamine, 25-mM HEPES, 5-ng/mL interleukin-2, and 1-μg/mL αCD28) in X-VIVO 15 (Lonza). T cells (2.5 × 10^5^) were seeded in 96-well plates previously coated with αCD3 (5 μg/ml in PBS, 200 μl/well, overnight, 4 °C). Cells were incubated for 72 h at 37 °C. After incubation, cells were transferred to 5 mL FACS tubes, washed once with PBS, resuspended in 50-μL FcX-SB (0.5 μL/test), and incubated for 10 min (4 °C). Next, 50 μL of Ab-SB-solution (0.5-μL/test CD4-PE, 1-μL/test CD8-FITC) were added to the cells and incubated for 30 min. After washing twice with SB, cells were fixed with IC fixation buffer (Thermo Fisher Scientific) for 10 min at 4 °C, centrifuged, and resuspended in 100 μL SB. Samples were measured on a FACS Canto II (BD Biosciences).

### Statistical analysis

Flow cytometric data were reported in cells/mg. Flow cytometric data compensation was performed using single stains. The “fluorescence-minus-one” strategy was used to calculate background fluorescence cutoffs (*54*). Data were analyzed using the FlowJo software (TreeStar). All statistical analyses were performed using GraphPad Prism 6.1 (GraphPad Software).

All variables were tested for Gaussian distribution using the Shapiro–Wilk normality test. Significant differences between two experimental groups with normal distribution were determined using unpaired student’s t-tests with Welch’
ss correction; otherwise the Mann–Whitney test was applied. To compare three or more groups, a one-way analysis of variance (ANOVA) was used, with Sidak’s post hoc test for multiple comparison corrections. Neutrophil content and MPO^+^ lymphocytes were correlated using Pearson’s correlation coefficient (r) and Spearman’s correlation coefficient rho (r_s_). The results were expressed as the mean and standard error of the mean, or standard deviation. A *p*-value of <0.05 was considered statistically significant, and it was denoted with one, two, three, or four asterisks when it was <0.05, 0.01, 0.001, or 0.0001, respectively.

## Supplementary Materials

Tables S1 and S2

Figures S1–S7

## Acknowledgments

We are grateful to Sabine Kern and Iris Red for their excellent technical assistance, as well as Sabine Donner for animal care. The experimental design diagrams were all created with BioRender.com.

## Funding

The lab of JK is supported by the OENB Anniversary Fund (17584), the FFG-Bridge 1 grant (871284), FWF-P35294, FWF, doctoral programs [PV-C, SR, AS DK-MOLIN (W1241), NCM (RespImmun, DOC-129), ZNM (DP-iDP, DOC-31)], OK (OEAW Doc Fellowship - 26477), and MK (BioTechMed). The lab work of RS is funded by the FWF grants P33325 and KLI887.

## Author contributions

Conceptualization and Supervision: PV-C, JK; Methodology and Investigation: PV-C, KM, NCM; Bioinformatics analysis: OK; Experiments: SR, MK, ZNM AS, AS, AL; Clinical database: LB, JL; Statistical analyses: PV-C, OK; Figures: PV-C; Data interpretation and Technical support: PV-C, OK, RS, WS, AH, GM, JK; Writing – original draft: PV-C and JK; Writing – review & editing: All authors.

## Competing interests

Authors declare that they have no competing interests.

## Notes

### Competing Interest Statement

The authors have declared no competing interest.

