## Supplementary Material for "Myeloperoxidase promotes a tumorigenic microenvironment in non-small cell lung cancer"

### Table of Contents

Tables S1 and S2

Figures S1-S7

**Table S1. Genes derived from the CPTAC-LUAD dataset.**

| MPO signature | | Control signature | | | |
|---|---|---|---|---|---|
| ADGRG3 | HIST1H4L | ACOXL | CLK4 | LRRC31 | TLR5 |
| ADM | IL1R2 | ADGRF5 | CNKSR2 | MAP3K15 | TMEM243 |
| AQP9 | LEP | ADHFE1 | CREBRF | MAPK10 | TNN |
| C11orf24 | LIF | ANK3 | CRYM | MR1 | WWC1 |
| CDH2 | LRFN4 | ANKDD1B | CYP4V2 | MS4A2 | ZDHHC15 |
| CEACAM3 | MRPL12 | ANKRD44 | CYSLTR1 | MYOZ1 | ZMAT1 |
| CEMIP | MTHFD2 | AR | ETFBKMT | N4BP2L1 | ZNF441 |
| CHD7 | NAMPT | ARHGAP31 | FAM161B | NFIX | ZNF540 |
| CTD-3088G3.8 | NFXL1 | ARHGDIB | FAM184A | NIPAL3 | ZNF554 |
| CXCL2 | NOP56 | ATP13A4 | FMO4 | NPC2 | ZNF563 |
| CXCL3 | P3H4 | ATP13A5 | FMO5 | NRN1L | ZNF763 |
| CXCL8 | PCBP1 | B3GAT1 | GAB1 | PIGR | ZNF846 |
| CYP24A1 | PI15 | C16orf89 | GANC | POU2F3 | ZRSR2 |
| ECT2 | PI3 | C17orf50 | GPR160 | PPP2R3A | |
| EIF4A1 | PLAUR | C1orf116 | GSAP | PTPN13 | |
| EIF5A | PLEKHG2 | CA1 | HAS3 | RBL2 | |
| ENTPD7 | PNO1 | CA13 | HLA-DPB1 | REPS2 | |
| FAM64A | PPIF | CACNA1F | HLF | RNASE1 | |
| FAM83D | PPM1G | CAMK2D | HNMT | RP11-490B18.9 | |
| FCAR | PVR | CAPN3 | HPGDS | SCTR | |
| FCGR3B | RCOR2 | CARF | HSD17B6 | SGF29 | |
| FKBP10 | SLC2A1 | CD1B | IL11RA | SLC15A2 | |
| FPR2 | SNRPA1 | CD1E | IL6R | SLC9A5 | |
| HCAR3 | SNRPD1 | CD302 | KAT2B | SLFN14 | |
| | TDG | CD40LG | KCNK5 | SMCO3 | |
| | TGM3 | CD74 | KIF13A | SOGA3 | |
| | TTL | CEBPA | KLHDC1 | TCEANC | |
| | UBE2S | CFAP221 | LANCL3 | TEF | |
| | UCK2 | CIRBP | LRP2BP | TLR3 | |

MPO Signature: genes that were significantly upregulated in the MPO high group.
Control Signature: genes that were significantly downregulated in the MPO low group.

**Table S2. Mouse and human flow cytometric antibody panels.**

| Panel | Antibody | Dilution | Clone | Company | Catalog number |
|---|---|---|---|---|---|
| **Tumor-infiltrating lymphoid immune cells (mouse)** | CD45-AF700 | 1:200 | 30-F11 | BioLegend | 103128 |
| | CD3-BUV395 | 1:40 | 145-2C11 | BD Biosciences | 563565 |
| | CD8-PerCPCy5.5 | 1:80 | 53-6.7 | BioLegend | 100734 |
| | CD4-BUV496 | 1:80 | GK1.5 | BD Biosciences | 564667 |
| | gdTCR-PECF594 | 1:40 | GL3 | BD Biosciences | 563532 |
| | PD-1-APC | 1:40 | 29F.1A12 | BioLegend | 135210 |
| | CD62L-BV605 | 1:50 | MEL-14 | BioLegend | 104438 |
| | CD44-BUV737 | 1:160 | IM7 | BD Biosciences | 612799 |
| | NKp46-BV510 | 1:20 | 29A1.4 | BioLegend | 137623 |
| | CD19-FITC | 1:160 | 6D5 | BioLegend | 115506 |
| | CD25-BV785 | 1:80 | PC61 | BioLegend | 102051 |
| | FoxP3-PE | 1:40 | FJK-16s | eBio | 12-5773-82 |
| **Tumor-infiltrating myeloid immune cells (mouse)** | CD45-BV785 | 1:160 | 30-F11 | BioLegend | 103149 |
| | Ly6C-APC | 1:79 | HK1.4 | BioLegend | 128015 |
| | Ly6G-PE /Dazzle | 1:166 | 1A8 | BioLegend | 127648 |
| | CD11c-BV605 | 1:20 | N418 | BioLegend | 117334 |
| | PD-L1-PeCy7 | 1:79 | 10F.9G2 | BioLegend | 124313 |
| | CD206-FITC | 1:160 | C068C2 | BioLegend | 141703 |
| | MHCII-PerCP-Cy5.5 | 1:160 | M5/114.15.2 | BioLegend | 107625 |

| | | | | | |
|---|---|---|---|---|---|
| | CD103-BV510 | 1:40 | 2E7 | BioLegend | 121423 |
| | CD11b-BUV737 | 1:80 | M1/70 | BD Biosciences | 612801 |
| | F4/80-BUV395 | 1:40 | T45-2342 | BD Biosciences | 565614 |
| | Siglec-F-PE | 1:40 | E50-2440 | BD Biosciences | 562068 |
| **Interferon-gamma expression (mouse)** | CD45-FITC | 1:100 | 30-F11 | BioLegend | 103108 |
| | CD3-BV421 | 1:25 | 145-2C11 | BioLegend | 100336 |
| | CD4-PE-Cy7 | 1:50 | RM4-5 | Biolegend | 100528 |
| | CD8-PerCP-Cy5.5 | 1:80 | 53-6.7 | BioLegend | 100734 |
| | IFN-γ-PE | 1:50 | XMG1.2 | BioLegend | 505808 |
| **T-cell proliferation (human)** | CD4- PE | 1:100 | RPA-T4 | BioLegend | 300508 |
| | CD8a-FITC | 1:100 | RPA-T8 | BioLegend | 301006 |
| **Interferon-gamma expression (human)** | CD4- PE-Cy7 | 1:25 | RPA-T4 | BioLegend | 300512 |
| | CD8-APC | 1:25 | RPA-T8 | BioLegend | 301014 |
| | IFN-γ-BUV395 | 1:20 | B27 | BD Biosciences | 563563 |
| **MPO binding/internalization (human)** | CD4-PE-Cy7 | 1:100 | RPA-T4 | BioLegend | 300512 |
| | CD8-BV510 | 1:50 | RPA-T8 | BioLegend | 301006 |
| | MPO-FITC | 1:2.5 | 5B8 | BD Biosciences | 340580 |
| **MPO in tumors from patients with NSCLC** | CD45-BV510 | 1:100 | H130 | BioLegend | 304035 |
| | CD66b-APC | 1:50 | G10F5 | BioLegend | 305118 |
| | CD163-BV605 | 1:25 | GH1/61 | BioLegend | 333616 |
| | MPO-FITC | 1:2,5 | 5B8 | BD Biosciences | 340580 |

**Fig. S1**

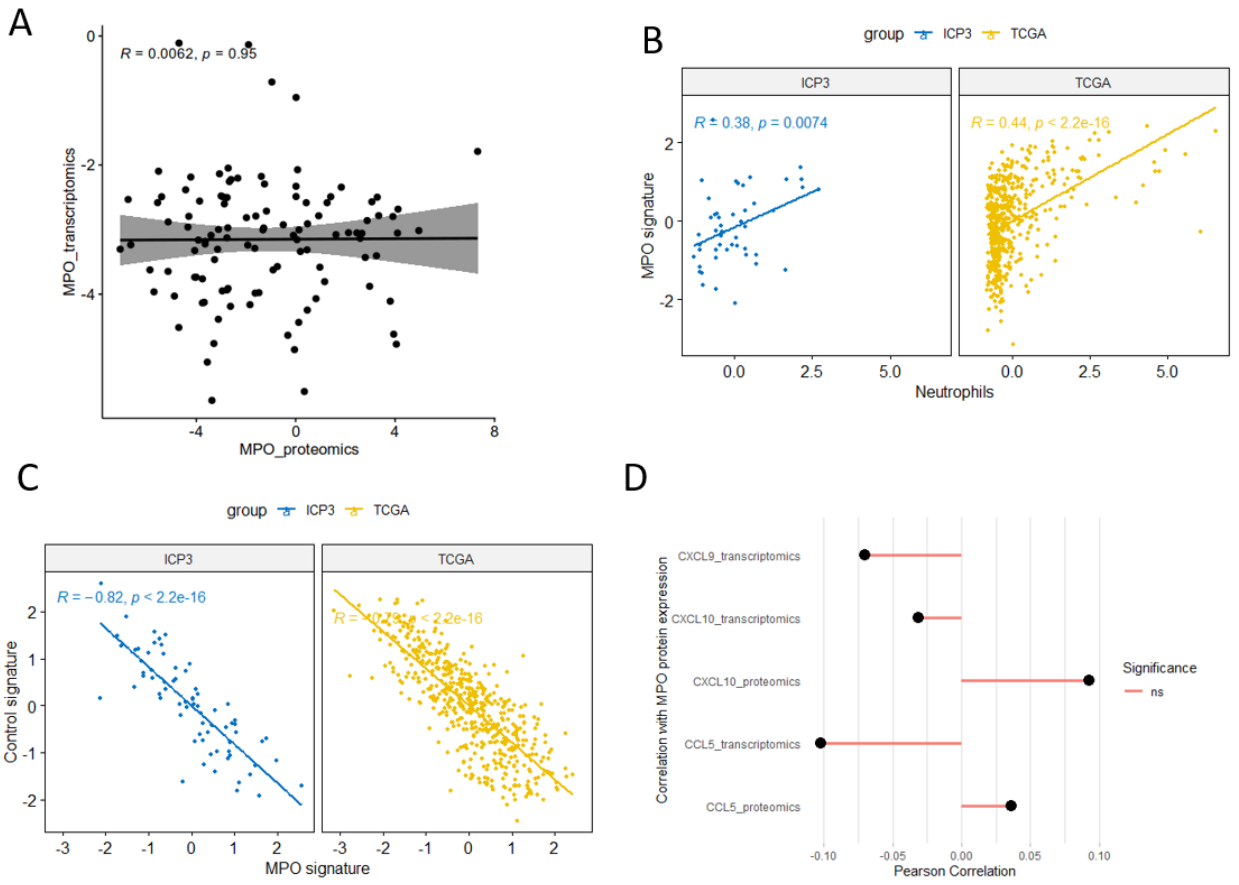


**Validation of MPO signature.**

**(A)** Scatter plot showing Pearson correlation between MPO transcript and protein expression in the CPTAC LUAD dataset. **(B)** Scatter plot showing Pearson correlation between scaled MPO signature and scaled Neutrophil content in two different datasets (ICP3: Neutrophils in % CD45+ cells; TCGA-LUAD: Percentage of Neutrophils derived from CASSANDRA website). **(C)** Scatter plot showing Pearson correlation between scaled MPO signature and Control signature in ICP3 and TCGA LUAD dataset. **(D)** Lollipop representation of Pearson correlation between selected cytokines and MPO protein expression.

**Fig. S2**

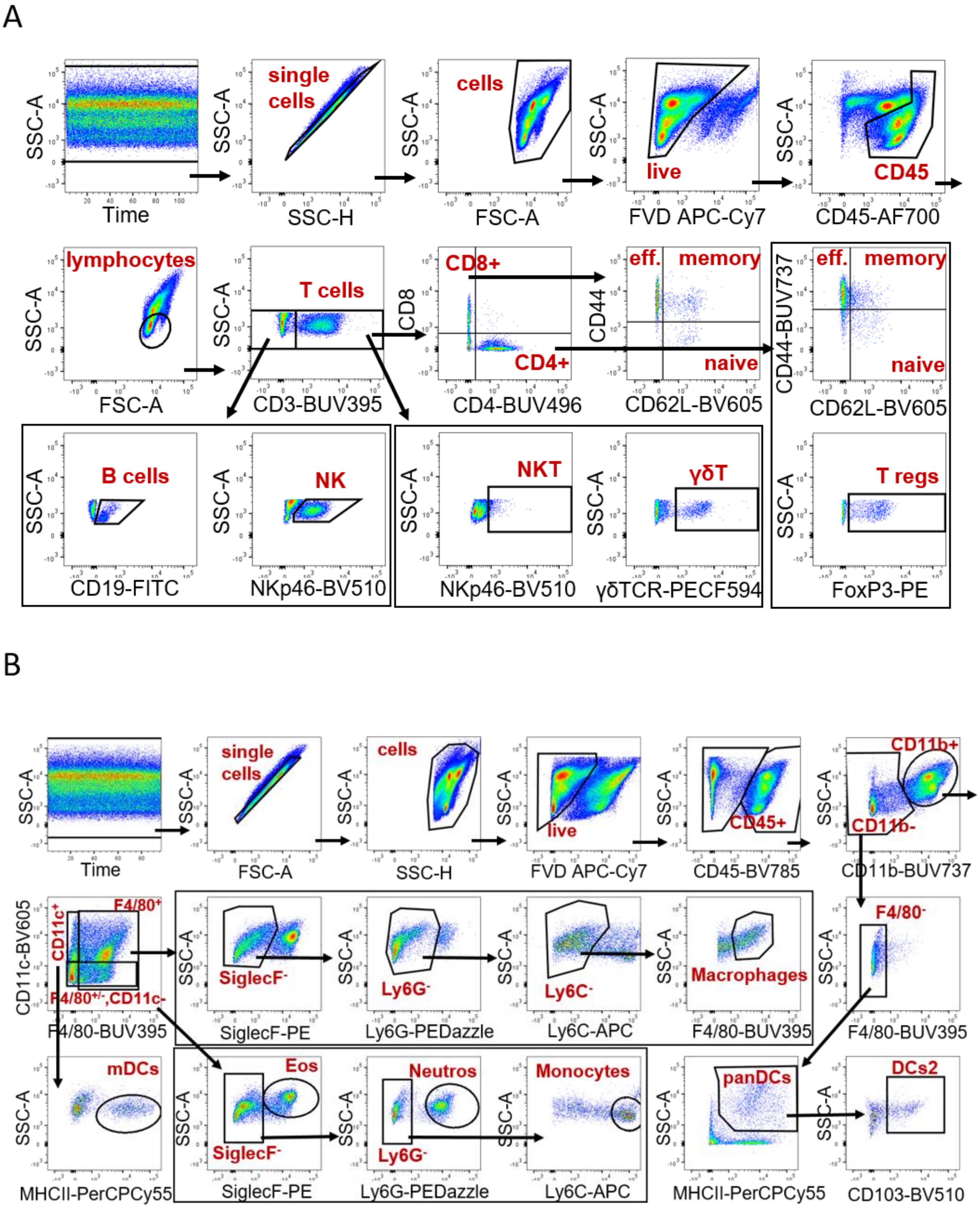
A
SSC-A
Time
single cells
SSC-H
cells
FSC-A
live
FVD APC-Cy7
CD45
CD45-AF700
lymphocytes
T cells
CD3-BUV395
CD8+
CD8
CD4+
CD4-BUV496
eff.
memory
naive
CD44
CD62L-BV605
CD44-BUV737
B cells
CD19-FITC
NK
NKp46-BV510
NKT
γδT
γδTCR-PECF594
T regs
FoxP3-PE
B
CD45+
CD45-BV785
CD11b+
CD11b-
CD11b-BUV737
CD11c-BV605
CD11c+
F4/80+
F4/80+/-,CD11c-
F4/80-BUV395
SiglecF-
SiglecF-PE
Ly6G-
Ly6G-PEDazzle
Ly6C-
Ly6C-APC
Macrophages
F4/80-
mDCs
MHCII-PerCPCy55
Eos
Neutros
Monocytes
panDCs
DCs2
CD103-BV510

C

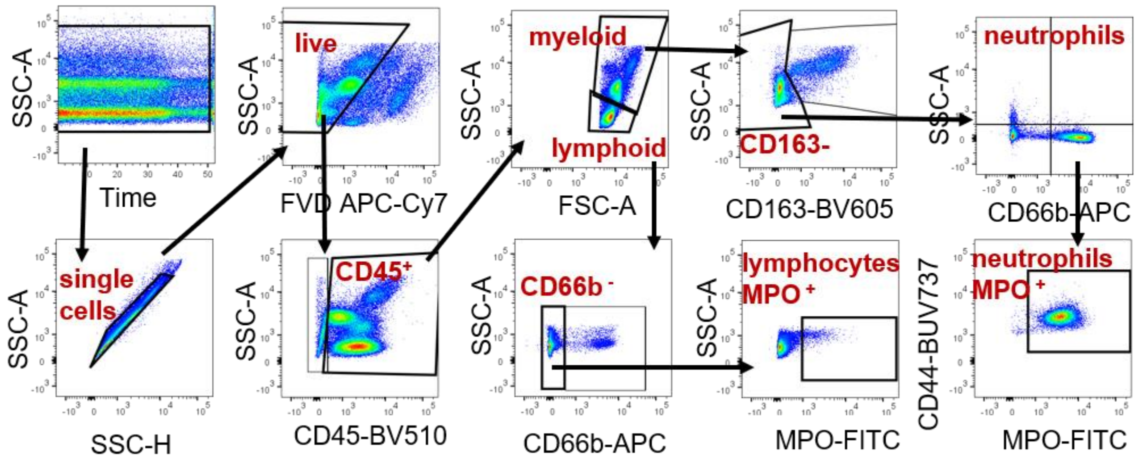


D

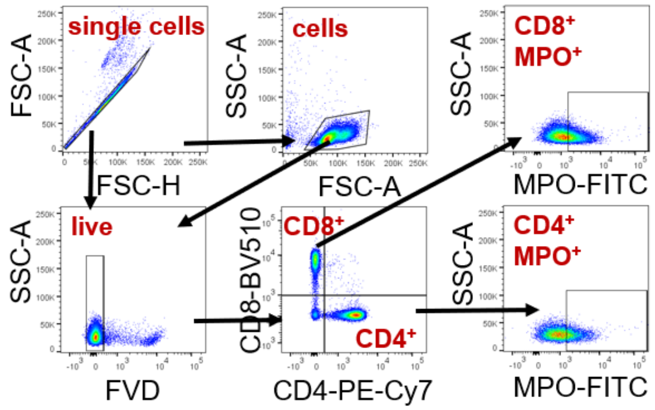


**Flow cytometric gating strategies.**

Schematic representation of the flow cytometric gating strategies used to analyze single cell suspensions obtained from KP and LLC cell tumors **(A and B)**, human lung tumors **(C)** and human lymphocytes from healthy donors **(D)**. Infiltrating CD45$^{+}$ cells were pre-gated for time and single cells. Dead cells were excluded, and an additional lymphocyte gate was used to exclude myeloid cells from the lymphoid panel.

**(A) Tumor-infiltrating lymphoid immune cells (mouse).** T cells were gated as CD45$^{+}$/CD3$^{+}$; NK cells as CD45$^{+}$/CD3$^{-}$/NKp46$^{+}$; B cells as CD45$^{+}$/CD3$^{-}$/CD19$^{+}$; γδ T cells as CD45$^{+}$/CD3$^{+}$/γδTCR$^{+}$; NKT cells as CD45$^{+}$/CD3$^{+}$/NKp46$^{+}$; CD8$^{+}$ T cells as CD45$^{+}$/CD3$^{+}$/CD8$^{+}$; CD4$^{+}$ T cells as CD45$^{+}$/CD3$^{+}$/CD4$^{+}$; CD8$^{+}$ T effector (eff.) cells as CD45$^{+}$/CD3$^{+}$/CD8$^{+}$/CD44$^{+}$; CD8$^{+}$ T naïve cells as CD45$^{+}$/CD3$^{+}$/CD8$^{+}$/CD62L$^{+}$; CD8$^{+}$ T memory cells as CD45$^{+}$/CD3$^{+}$/CD8$^{+}$/CD44$^{+}$/CD62L$^{+}$; CD4$^{+}$ T effector (eff.) cells as CD45$^{+}$/CD3$^{+}$/CD4$^{+}$/CD44$^{+}$; CD4$^{+}$ T naïve cells as CD45$^{+}$/CD3$^{+}$/CD4$^{+}$/CD62L$^{+}$; CD4$^{+}$ T memory cells as CD45$^{+}$/CD3$^{+}$/CD4$^{+}$/CD44$^{+}$/CD62L$^{+}$; and CD4$^{+}$ T regulatory (T regs) as CD45$^{+}$/CD3$^{+}$/CD4$^{+}$/FoxP3$^{+}$. PD-1 expression was as percentage of CD4$^{+}$ and CD8$^{+}$ T cells.

**(B) Tumor-infiltrating myeloid immune cells (mouse).** Eosinophils were gated as $CD45^+/CD11b^+/CD11c^-$/Siglec-$F^+$; neutrophils as $CD45^+/CD11b^+/CD11c^-$/Siglec-$F^-$/$Ly6G^+$; monocytes as $CD45^+/CD11b^+/CD11c^-$/Siglec-$F^-$/$Ly6G^-/Ly6C^+$ and macrophages as $CD45^+/CD11b^+/CD11c^{-/+}$/Siglec-$F^-$/$Ly6G^-/Ly6C^-/F4/80^+$; myeloid dendritic cells (mDCs) as $CD45^+/CD11b^+$/ $F4/80^{+/-}/CD11c^+/MHCII^+$; plasmacytoid DCs (panDCs) as $CD45^+/CD11b^-$/ $F4/80^-/MHCII^+$ and dendritic cells (DCs2) $CD45^+/CD11b^-$/ $F4/80^-/MHCII^+/CD103^+$.

**(C) MPO in tumors from patients with NSCLC**. CD66b marker was used to exclude myeloid cells within the lymphoid gate and MPO presence in lymphocytes was determined as $CD66b^-$ /$MPO^+$. $MPO^+$ neutrophils were gated as $CD163^-/CD66b^+/MPO^+$.

**(D) MPO binding/internalization (human).** MPO presence in $CD4^+$ and $CD8^+$ T cells was determined as $CD4^+/MPO^+$ and $CD8^+/MPO^+$, respectively.

**Fig. S3**

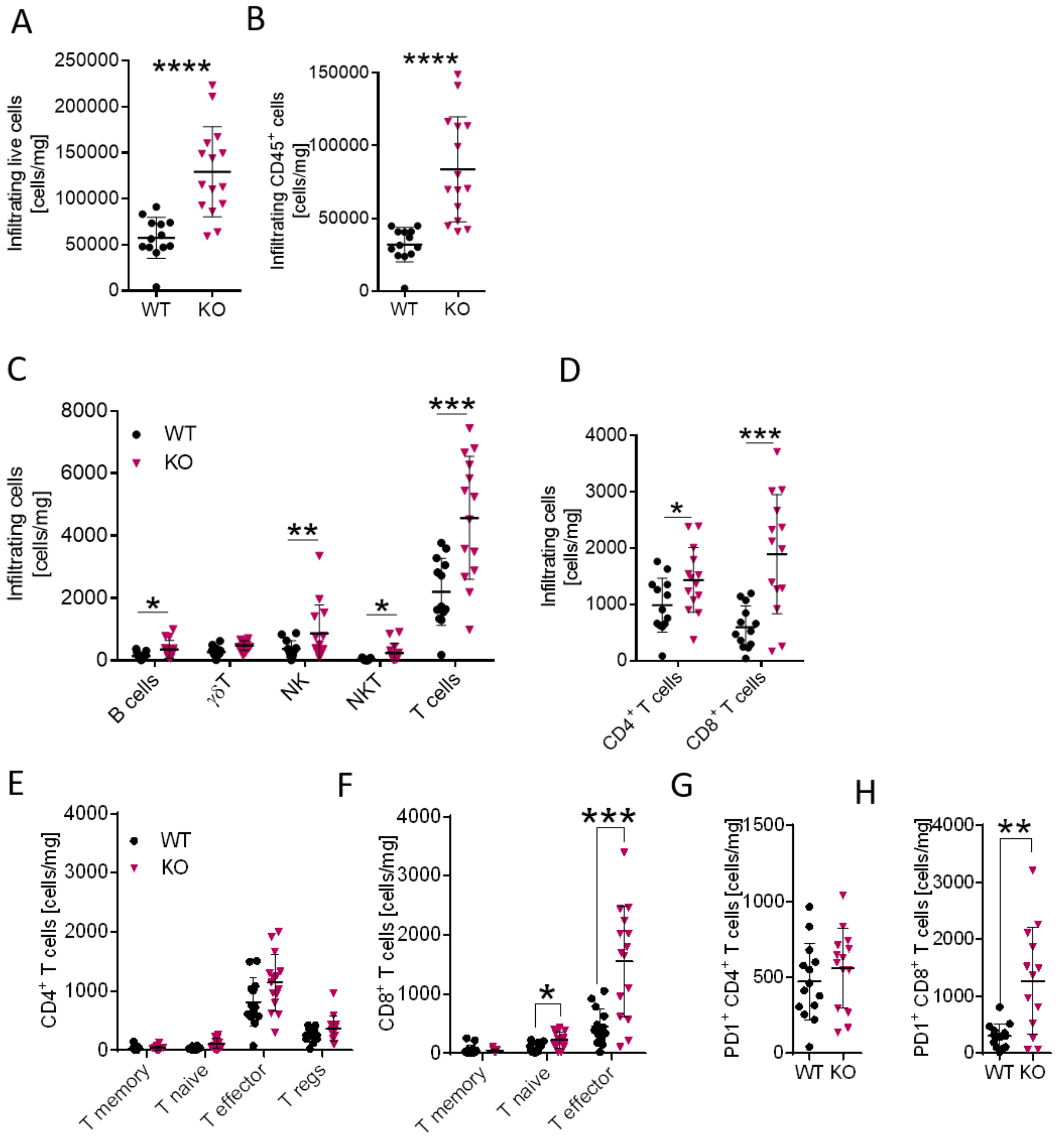


**Infiltrating immune cells in LLC tumors.**

**(A–H)** Flow cytometric analysis of single-cell suspensions of LLC tumors from MPO wild-type (WT) and knockout (KO) mice. The data were pooled from two independent experiments ($n$ = 15–16) and expressed as means ± standard deviations. All variables were tested for Gaussian distribution using the Shapiro–Wilk normality test. Statistical differences between WT and KO mouse data with normal distribution were determined using unpaired student's t-tests with Welch's correction; otherwise, the Mann–Whitney test was applied (*$p$ < 0.05, **$p$ < 0.01 and ***$p$ < 0.001).

**Fig. S4**

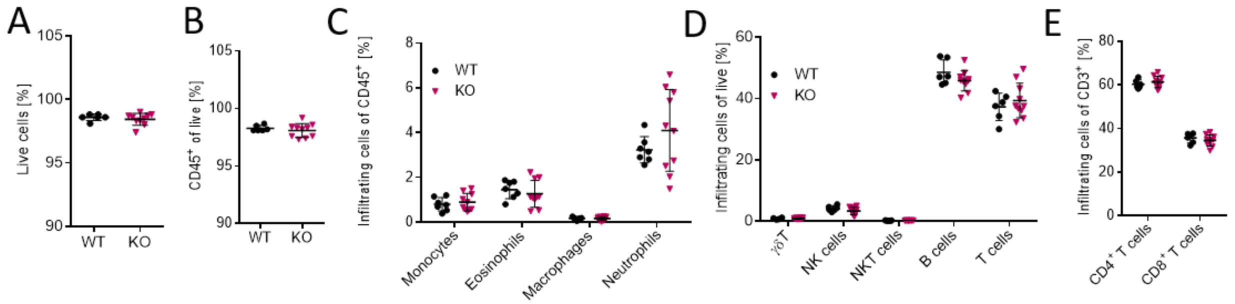


**Splenic controls of tumor-free mice.**

**(A-E)** Flow cytometric analysis showing immune cells content on spleens from tumor-free mice. The data were obtained from one independent experiment (n = 6–10) and expressed as means ± standard deviations. Statistical differences between WT and KO mouse data were determined using unpaired student's t-tests with Welch's correction. No significant differences were found in any of the tested groups.

**Fig. S5**

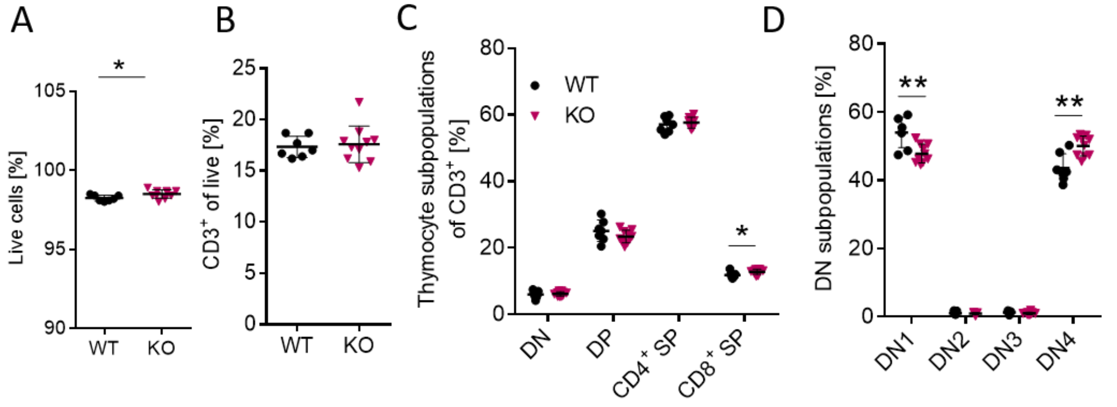


**Thymic controls of tumor-free mice.**

**(A-E)** Flow cytometric analysis showing immune cells content on thymus from tumor-free mice. The data were obtained from one independent experiment (n = 7–10) and expressed as means ± standard deviations. Statistical differences between WT and KO mouse data were determined using unpaired student's t-tests with Welch's correction (*$p < 0.05$ and **$p < 0.01$).

**Fig. S6**

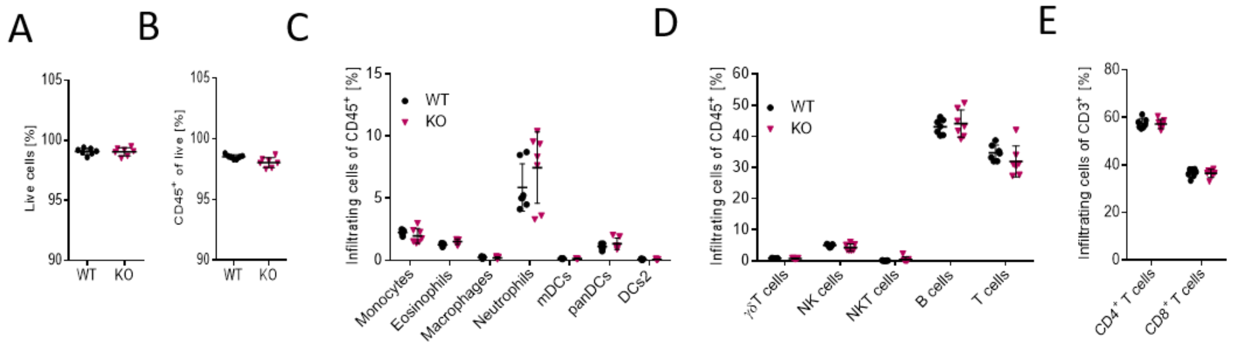


**Splenic controls of KP tumor-bearing mice.**

**(A-E)** Flow cytometric analysis showing immune cells content on spleens from tumor-free mice. The data were obtained from one independent experiment (n = 7) and expressed as means ± standard deviations. Statistical differences between WT and KO mouse data were determined using unpaired student's t-tests with Welch's correction. No significant differences were found in any of the tested groups.

**Fig. S7**

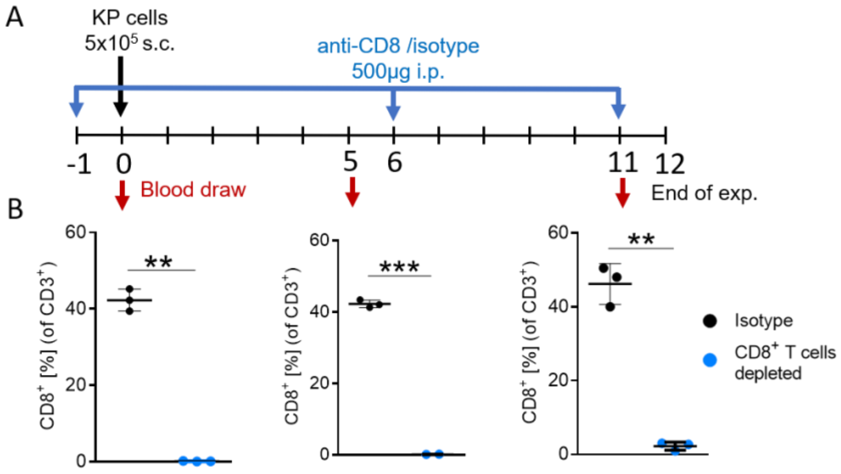


**CD8$^{+}$ T cells depletion controls.**

**(A)** 500 µg of anti-CD8 antibody or isotype was intraperitoneally (i.p.) administered one day prior to the subcutaneous (s.c.) engraftment of $5 \times 10^{5}$ KP cells into $MPO^{+/+}$ (WT) and $MPO^{-/-}$ (KO) mice. 500 µg and 250 µg of anti-CD8 antibody or isotype were administered in days 5 and 11, respectively. Blood was drawn on days 0, 6 and 11. **(B)** Expression of CD8$^{+}$ T cells was determined by flow cytometry. Statistical differences between WT and KO with normal distribution were determined using unpaired student's t-test with Welch´s correction (**p<0.01, ***p<.001).
